# An Enhanced Pipeline for Multi-Omic Integration Based on Topological Data Analysis

**DOI:** 10.64898/2026.09.21.752955

**Authors:** Veronica Paparozzi, Anna Plaksienko, Marco Pedicini, Christine Nardini

## Abstract

The advent of high-throughput sequencing technologies has made it essential to employ advanced tools for data integration and interpretation. In this work, we propose to build upon two existing efforts by expanding their scope and applicability to multi-omic data integration through the development of a pipeline that generates a topologically informed ranking of genes to assess gene relevance to disease. Specifically, we exploit topological data analysis, capitalizing on: its graph-based framework, to refine complex interactions characteristic of biological systems; its intrinsic explainability, to enable interpretation of model outputs in terms of biologically meaningful features; its robustness to small input perturbations, to ensure stable results in the presence of noisy data, as omics are. In particular, we employ Harmonic Persistent Homology (HPH), and propose two relevant advances. First, we adopt an integration and clustering tool (i.e. iNETgrate) as an informed means to reduce data size and integrate multiple omic layers by aggregating methylation beta values from loci to the gene level, to be combined with gene expression values. This enables the application of HPH, otherwise limited by the computational burden. Second, as a non-trivial consequence, we apply HPH to multi-omic molecular data, and not to patients as done so far, enlarging HPH scope. Finally, we follow up on recent efforts toward performance standardization, and validate our results against biomarkers from a well-known TCGA breast cancer benchmark dataset.

## 1 Introduction

Advances in high-throughput technologies have enabled the comprehensive profiling of biological systems at multiple molecular levels, including the genome, transcriptome, proteome, metabolome and epigenome, to name a few. These data layers provide a complementary and interconnected view to describe a biological system. The analysis of multi-omic data, however, poses significant challenges due to different scales and representations (e.g., gene- versus locus-level), thus, the inherent biologic complexity can be captured by different approaches, leading to multiple methodologies, with pros and cons [1].

A popular direction assumes that data are affected by latent variables that cannot be measured directly (e.g. underlying biological processes). Methods such as Multi-Omics Factor Analysis [2] discover shared sources of variation of multiple omic layers through matrix factorization. The identified factors can be used for downstream analyses like identification of outliers, sample subgroups and data imputation. Another direction exploits supervised classification using deep learning: MOGONET (Multi-Omics Graph cOnvolutional NETworks) [3] jointly models omic-specific patterns and cross-omic label correlations for each layer before integration. Tools such as Similarity Network Fusion [4] produce networks where nodes represent patients and edges represent similarity between them based on all available omic layers.

Focusing specifically on the integration of transcriptomics and methylomics, iNETgrate [5] appears to be an ideal tool that constructs a graph with genes as vertices and edges weighted by corresponding transcriptomics and DNA methylation values simultaneously. Its main advantages lie in the ability to handle the one-to-many relationship between genes and CpG sites in a reasonable time frame, to produce modules collecting similar genes. Yet, modules are of varying size and interpretation of large ones (hundreds of molecules) is not straightforward. To overcome this limitation, we sought to integrate the analysis of such modules with Topological Data Analysis (TDA). TDA is a framework for characterizing the shape of data using tools from algebraic topology, with Persistent Homology (PH) as a central method [6]. PH provides an unsupervised and interpretable approach that captures higher-order, network-like relationships, making it well suited for biological data. Importantly, its stability with respect to distances between PH outputs ensures robustness to noise, a key property for multi-omic applications. Harmonic Persistent Homology (HPH) [7] extends PH by further enhancing interpretability, and it has recently been used for the first time in application to omics and multi-omics [8]. However, although transcriptomic applications were used to identify genes signatures, multiomic applications were patients-centered. The HPH approach, in fact, presents scalability limits, which coupled to the curse of dimensionality accompanying omics, makes molecular multi-omic signature identification challenging. We therefore propose a pipeline to expand to multi-omic applications by combining first the omics’ integration and clustering method to harmonize methylation with transcription data while reducing the dimensionality (i.e. iNETgrate), and then apply HPH (i.e. maTilDA [8]) to derive gene-level insights by investigating harmonic weights for assessing relevance. This pipeline has been tested on a known dataset [9] with conventional ranking metrics against a standard benchmark [10] and with an original HPH score in comparison to single omic analyses.

## 2 Data and Methods

### 2.1 Data

Transcriptomics and methylomics are publicly available at https://gdc.cancer.gov/about-data/publications/brca_2012[9]. Samples were filtered to retain the tumor samples from patients with both transcriptomic and methylomic data. After initial pre-processing, we were left with 266 samples in the transcriptomic layer (17 814 genes) and 224 samples in the methylation layer (323 690 probes matched to at least one gene). Samples were matched across omic layers by patient and sample IDs. Biomarkers are available at https://zenodo.org/records/17993142 as *drug bks.csv*.

### 2.2 Data integration and selection

We used the `iNETgrate` R package [5] to integrate gene expression and methylation beta values. Briefly, in the network we aim to estimate, every node represents a gene and its corresponding CpG sites. First, methylation levels across multiple CpG sites are reduced to one number (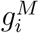 for gene *i*). If the number of loci corresponding to a gene is less than six, weighted average of *β*-values is calculated using PCA. Otherwise, iNETgrate first identifies the most connected cluster of loci for each gene, and then computes the aggregated value based only on that subset. Afterwards, a weighted network is constructed with *integrated similarity* values based on absolute correlation of gene expression (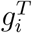 for gene *i*) and DNA methylation levels (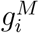 for gene *i*) for each pair *i, j* of genes as

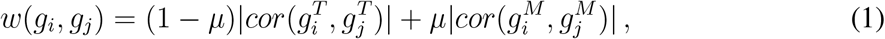

with *μ* controlling the relative contribution of each layer. We built three networks, using *μ* = 0.6 (MT), following iNETgrate vignette, as well as *μ* = 0 (network based solely on gene expression, T) and *μ* = 1 (network based solely on methylation, M). After the network is constructed, iNETgrate performs hierarchical clustering to identify modules (groups) of genes that exhibit similar patterns of expression and methylation (topological overlap is used).

### 2.3 Signatures identification

A standard TDA pipeline begins by representing data as an *abstract simplicial complex* whose elements are called *simplices*. In our setting, 0-simplices (i.e. nodes) correspond to genes and 1-simplices (i.e. edges) encode for correlations between the omic data as output by iNETgrate (see Equation 1 and details on distance matrix below). PH then tracks the evolution of topological features (i.e., *holes*), such as connected components (0-dimensional), loops (1-dimensional), and voids (2-dimensional), across a filtration parameter, here correlation values. Intuitively, PH describes how the graph of each iNETgrate module topologically evolves as edges are progressively added during the filtration. This is represented in a *barcode*, where the y-axis records the features and the x-axis the filtering parameter (distance). The length of the bar is a proxy for the stability of the corresponding feature. Harmonic Persistent Homology (HPH) [7] assigns a unique *harmonic cycle* to each bar, obtained by weighting simplices according to their contribution across all cycles associated with the given feature. The resulting harmonic weights can then be propagated to vertices (genes), yielding node-specific importance scores for each bar, a.k.a. *HPH weights*. For clustering computation, genes are represented as vectors of HPH weights, which, by construction, better capture the similarity among molecules.

The `maTilDA` (https://github.com/IBM/matilda) [8] tool constructs simplicial complexes using as input distance matrices, then computes PH and harmonic cycles. It returns a dataframe of harmonic weights with one row per bar and one column per node (i.e. molecule). We applied `maTilDA` to perform TDA on each distinct module produced by iNETgrate considering the distance matrix values defined as *d*_*ij*_ = 1 *−w*(*g*_*i*_, *g*_*j*_) for all *i, j* in the module.

Parameters were set compromising between data retention and computational cost: (i) simplicial complexes with *d*_*ij*_ *<* 0.75 for all *i, j* (threshold higher than the sum of the mean and standard deviation of all modules), yielding 1-complexes; (ii) PH bars length set to limit the number of projected representative cycles to 9000. To extract clusters, we used SciPy’s hierarchical clustering, with cosine distance and single linkage applied to the dataframe columns (vectors). Finally, for each gene we compute the *HPH score* as the sum of its HPH weights normalized over the number of 1-holes (1-bars) in the module. By construction, this score encodes the relevance of a node within the module’s features, and therefore represents a proxy for the relevance of the gene in the analysis; normalization mitigates the increased number of 1-holes present in larger modules.

### 2.4 Performances against benchmark

To validate the results, we used a recently published effort that provides multi-omic standards [10] with a curated sets of biomarkers supported by clinical evidence on TCGA broadly used datasets, and a list of standard metrics. In particular we used the drug-related biomarkers set: 94 manually curated molecules for breast cancer, and the single score metrics (optimal value 1): Average Recall (AR), Normalized Discounted Cumulative Gain (NDCG), Reciprocal Rank (RR), Area Under the Receiver Operating Characteristic Curve (AUROC), and one-sided Mann–Whitney U test (MW) on the ranking positions of biomarkers versus non-biomarkers. Lower *p*-values in MW provide stronger evidence that biomarkers rank higher than non-biomarkers. Stability measures were not included, as our method does not use training sets.

### 2.5 Performances against single omics

To gain a more general view on the opportunity to integrate multi-omics we performed a comparison among the three iNETgrate cases T, M, MT. Since the MT, T and M cases lead to different, non overlapping modules, we operated at the *global* level, i.e. we sorted all the HPH scores to rank genes irrespective of their original module. We then identified the number of biomarkers whose rank is improved (Number of Positive Differences, NPD) or impaired (Number of Negative Differences, NND) in the MT case versus the T or M case. Finally, we also quantified these differences in terms of differential HPH scores, namely the Sum of Positive Differences (SPD) and the Sum of Negative Differences (SND), respectively, defined for *X* ∈ {*M, T*} as:

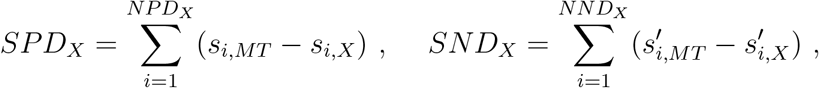

where 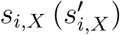 denotes the HPH score of the biomarker relative to the *i*-th positive (resp. negative) difference in the *X* case, and *·*_*X*_ denotes a measure w.r.t. the MT vs X comparison.

## 3 Results

### 3.1 Data integration and selection

`iNETgrate` MT analysis identified 166 modules ranging in size from 1320 to 5 genes (default smallest size for hierarchical clustering). Gene network based solely on gene expression (T) had 41 modules (largest 1197 genes). The network based on DNA methylation (M), consisted of 105 modules (largest 1631 genes).

### 3.2 Signatures identification and performances against benchmark

The MT network contains 38 of the 94 drug-related biomarkers (26 in T and 48 in M). Performance metrics for MT modules containing at least one biomarker are shown in Table 1. Among those, Module 7 (251 genes) includes three biomarkers (AGR2, CCND1 and ESR1) and presents the most favorable performance metrics among modules containing more than one biomarker, including a significant MW *p*-value. Therefore, we focus our attention on this module. The heatmap of Module 7 harmonic weights clearly visually identifies a cluster (that corresponds to cosine similarity 0.75) of 13 genes all included in the top-25 HPH scores, and highlighted in red in Figure 1. Performances were also computed across the (0.6–0.85) cosine similarity range, with minor differences in 0.68-0.75.

**Table 1:**
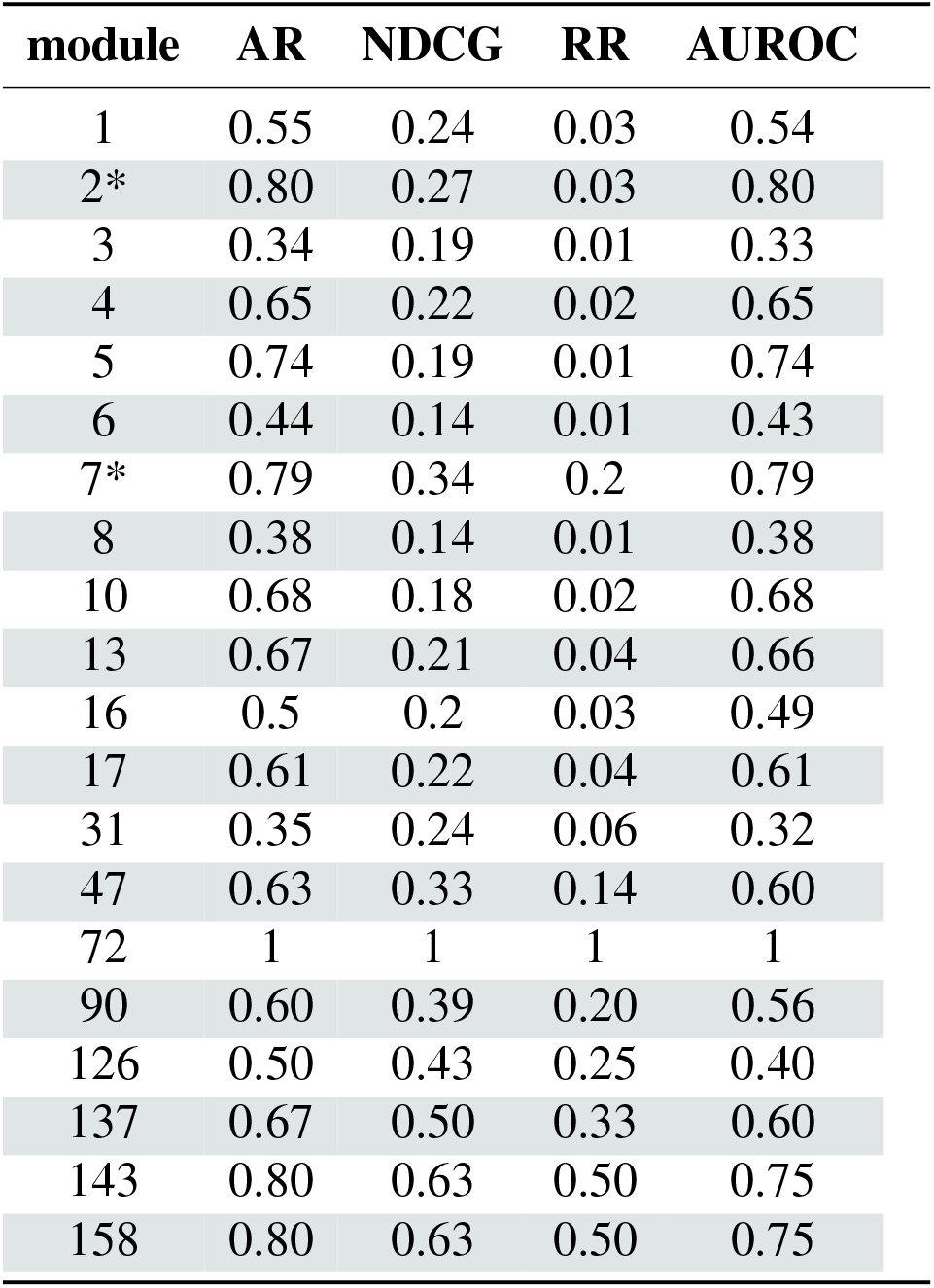
Results of performance metrics for gene ranking in the MT case. Metrics are computed for each module detected by iNETgrate using drug-related biomarkers. MW *p*-values are indicated as significance levels (* *p <* 0.05)

| module | AR | NDCG | RR | AUROC |
| --- | --- | --- | --- | --- |
| 1 | 0.55 | 0.24 | 0.03 | 0.54 |
| 2* | 0.80 | 0.27 | 0.03 | 0.80 |
| 3 | 0.34 | 0.19 | 0.01 | 0.33 |
| 4 | 0.65 | 0.22 | 0.02 | 0.65 |
| 5 | 0.74 | 0.19 | 0.01 | 0.74 |
| 6 | 0.44 | 0.14 | 0.01 | 0.43 |
| 7* | 0.79 | 0.34 | 0.2 | 0.79 |
| 8 | 0.38 | 0.14 | 0.01 | 0.38 |
| 10 | 0.68 | 0.18 | 0.02 | 0.68 |
| 13 | 0.67 | 0.21 | 0.04 | 0.66 |
| 16 | 0.5 | 0.2 | 0.03 | 0.49 |
| 17 | 0.61 | 0.22 | 0.04 | 0.61 |
| 31 | 0.35 | 0.24 | 0.06 | 0.32 |
| 47 | 0.63 | 0.33 | 0.14 | 0.60 |
| 72 | 1 | 1 | 1 | 1 |
| 90 | 0.60 | 0.39 | 0.20 | 0.56 |
| 126 | 0.50 | 0.43 | 0.25 | 0.40 |
| 137 | 0.67 | 0.50 | 0.33 | 0.60 |
| 143 | 0.80 | 0.63 | 0.50 | 0.75 |
| 158 | 0.80 | 0.63 | 0.50 | 0.75 |

**Figure 1:**
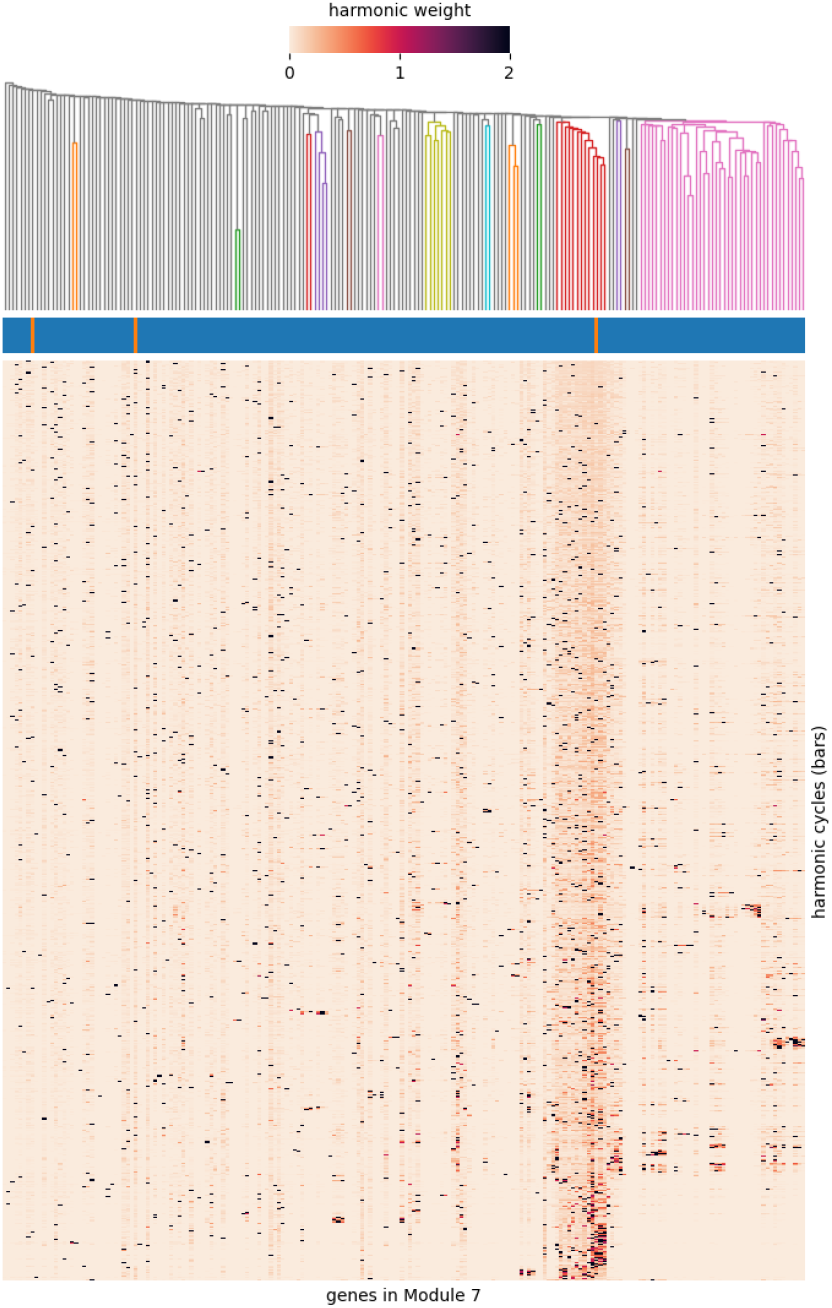
Dendrogram and heatmap based on node-level weights in Module 7. Bottom: heatmap of node-level weights (bars longer than 0.3). Middle: orange rectangles indicate biomarkers. Top: dendrogram (cosine similarity) cut at 0.75.

### 3.3 Single-vs multi-omic global gene ranking

To evaluate the contribution of the multi-omic integration we computed the global ranking under the three cases MT, T and M, shown in Table 2. First we can observe that all performance metrics, except for RR which considers only the first correct rank, are improved or at least similar to those obtained when considering a single omic layer, confirming the idea that multi-omic integration carries more information.

**Table 2:** Global ranking comparison. Left: Global ranking results for the MT, M, and T cases. MW *p*-values are indicated as significance levels (*** *p <* 0.001). Right: Comparison of positive and negative differences between the MT versus M and T cases. NPD: Number of Positive Differences; NND: Number of Negative Differences; SPD: Sum of Positive Differences; SND: Sum of Negative Differences.

| case | AR | NDCG | RR | AUROC | case | NPD | NND | SPD | SND |
| --- | --- | --- | --- | --- | --- | --- | --- | --- | --- |
| MT*** | 0.69 | 0.47 | 0.33 | 0.70 |  |  |  |  |  |
| M*** | 0.63 | 0.43 | 0.50 | 0.63 | MT vs M | 28 | 9 | 6.65 | -0.15 |
| T*** | 0.70 | 0.41 | 0.33 | 0.71 | MT vs T | 10 | 11 | 2.96 | -0.31 |

To further deepen the observation and perform the comparison focused on the biomarkers HPH scores, Table2 (right panel) compares single- and multi-omic cases at the biomarker level.

Notably, although the number of improved biomarkers in the global rankings of “MT vs T” is slightly lower (*NPD* = 10) than that of impaired ones (*NND* = 11), the aggregate magnitude of the positive differences of HPH scores (*SPD* = 2.96) is greater than that of the negative ones (*SND* = *−*0.31), suggesting that HPH scores are able to capture relevant information.

## 4 Conclusion

In this work, we explored the possibility to retain the best properties of two approaches for the integrated analysis of omic data, while overcoming their limitations. Given our specific interest in methylomics and transcriptomics we started from iNETgrate, able to manage the association between CpG and transcripts, and refined the large output module with a TDA-based approach. HPH enabled the identification of a robust and very manageable final molecular signature in a well known benchmark dataset across numerous metrics with the resulting rankings exhibiting consistency with the current state of biomarker knowledge, as supported by the evaluation metrics. The global ranking comparison further suggests that integrating multiple omics in this analysis can yield better overall performance than analyzing individual omics layers separately. Several limitations have to be highlighted. First, comparison with other multi-omic integration methods, such as above-mentioned MOFA, SNF and others, was out of scope for this short manuscript, and will be included in future work. Then, due to the computational cost of the TDA analysis, we limited our study to the more relevant, longest persistence bars (9000), since shorter bars are often interpreted as noise. Nevertheless, future work should more carefully test this hypothesis. Future work will additionally include a more systematic validation on additional benchmark data, and the ambition to apply it to experiments of our direct interest.

## Acknowledgments

V.P. is a Ph.D. student enrolled in the National Ph.D. in AI, course on Health and Life Sciences, organized by Università Campus Bio-Medico di Roma in collaboration with IAC-CNR.

## Data availability

The raw breast cancer data used in this manuscript are publicly available at https://gdc.cancer.gov/about-data/publications/brca_2012 as *BRCA*.*Gene Expression*.*Level 3*.*tar* (microarray gene expression) and *BRCA*.*HumanMethylation450*.*Level 3* (microarray methylation levels). The biomarkers supported by clinical evidence are available at https://zenodo.org/records/17993142 as *drug bks.csv* in the *data*.*zip* file.

